# The Ceratodon purpureus genome uncovers structurally complex, gene rich sex chromosomes

**DOI:** 10.1101/2020.07.03.163634

**Authors:** Sarah B. Carey, Jerry Jenkins, John T. Lovell, Florian Maumus, Avinash Sreedasyam, Adam C. Payton, Shenqiang Shu, George P. Tiley, Noe Fernandez-Pozo, Kerrie Barry, Cindy Chen, Mei Wang, Anna Lipzen, Chris Daum, Christopher A. Saski, Jordan C. McBreen, Roth E. Conrad, Leslie M. Kollar, Sanna Olsson, Sanna Huttunen, Jacob B. Landis, J. Gordon Burleigh, Norman J. Wickett, Matthew G. Johnson, Stefan A. Rensing, Jane Grimwood, Jeremy Schmutz, Stuart F. McDaniel

**Author notes:** Department of Crop, Soil, and Environmental Sciences, Auburn University, Auburn, AL, USA. HudsonAlpha Institute for Biotechnology, Huntsville, AL, USA.

## Abstract

Non-recombining sex chromosomes, like the mammalian Y, often lose genes and accumulate transposable elements, a process termed degeneration^1,2^. The correlation between suppressed recombination and degeneration is clear in animal XY systems^1,2^, but the absence of recombination is confounded with other asymmetries between the X and Y. In contrast, UV sex chromosomes, like those found in bryophytes, experience symmetrical population genetic conditions^3,4^. Here we test for degeneration in the bryophyte UV sex chromosome system through genomic comparisons with new female and male chromosome-scale reference genomes of the moss *Ceratodon purpureus*. We show that the moss sex chromosomes evolved over 300 million years ago and expanded via two chromosomal fusions. Although the sex chromosomes show signs of weaker purifying selection than autosomes, we find suppressed recombination alone is insufficient to drive gene loss on sex-specific chromosomes. Instead, the U and V sex chromosomes harbor thousands of broadly-expressed genes, including numerous key regulators of sexual development across land plants.

## Main text

Sex chromosomes arise when an ordinary pair of autosomes gains the capacity to determine sex^5^. A defining characteristic of sex chromosomes is suppressed recombination in the heterogametic sex. It is widely believed this lack of meiotic recombination makes natural selection less effective, predisposing non-recombining chromosomes, like the mammalian Y, to degeneration and gene-loss^1,2^. However, although some non-recombining chromosomes rapidly degenerate, or are completely lost, the sex chromosomes in other groups remain homomorphic or expand^1^. This diversity of form and gene content suggests the role of suppressed recombination in the long-term trajectory of sex chromosome evolution must be modulated by other processes related to the life history of the organism. Identifying these important processes requires comparative analyses across multiple eukaryotic lineages.

Many organisms, including bryophytes, algae, and some fungi, have a haploid UV sex chromosome system, in which females inherit a non-recombining U and males inherit a non-recombining V^3,4^. The sex-specific transmission pattern of both chromosomes means factors that are confounded in XY or ZW systems, such as suppressed recombination, hemizygosity, and sex-limited inheritance, are independent on UV chromosomes. Many UV sex chromosome systems may be ancient^4^, providing ample time for degenerative processes to act. However, the structural complexity of sex chromosomes has precluded genomic analyses in UV systems. Here we evaluate the relative roles of gene gain and degeneration in shaping the evolution of the bryophyte UV sex chromosomes using new chromosome-scale female and male genomes of the moss, *Ceratodon purpureus*.

## Results

Ancestral-state reconstructions of dioecy suggest that sex chromosomes evolved early in the history of the extant mosses^6^. To reconstruct the evolutionary history of the bryophyte UV sex chromosomes, we assembled and annotated chromosome-scale genomes of the GG1 (female) and R40 (male) *C. purpureus* isolates. Although the *C. purpureus* genome is relatively small, the sex chromosomes are large and have extensive repeat content, making them a challenge to assemble ^7^, particularly with short-read technologies, which often do not span a whole repeat. We therefore used a combination of Illumina, Bacterial Artificial Chromosomes (BACs), PacBio, and Dovetail Hi-C (Supplementary Tables 1-4; Extended Data Fig. 1). The version 1.0 genome assembly of R40 comprises 358 Megabases (Mb) in 601 contigs (N50 1.4 Mb), with 98.3% of the assembled sequence in the largest 13 pseudomolecules, corresponding to the 13 chromosomes in its karyotype^8^. The version 1.0 GG1 assembly is 349.5 Mb in 558 contigs (N50 1.4 Mb), with 97.9% of assembled sequence in the largest 13 pseudomolecules. Using over 1.5 billion RNA-seq reads for each of the genome lines and additional *de novo* assemblies of other *C. purpureus* isolates (Supplementary Table 5), we annotated 31,482 genes on the R40 assembly and 30,425 on GG1 (BUSCO v3.0 of 69% using Embryophyte; 96.7 and 96.4%, respectively using Eukaryote; values similar to the moss *Physcomitrium patens*^*9*^).

To examine the conservation of genome architecture, we performed synteny analyses between the two *C. purpureus* genomes and the *P. patens* genome. GG1 and R40 were collected from distant localities (Gross Gerungs, Austria and Rensselaer, New York, USA, respectively)^10^, and we found the assemblies had numerous structural differences (Fig. 1). In the self-synteny analysis we found clear homeologous chromosome pairs resulting from an ancient whole-genome duplication (Fig. 1, Extended Data Fig. 1), consistent with previous transcriptomic^10,11^ and our own *Ks*-based analyses (Extended Data Fig. 1; Supplementary Table 6). We also identified abundant synteny between the *C. purpureus* and *P. patens* chromosomes, which diverged over 200 million years ago (MYA)^12^ (Fig. 1). This result demonstrates the ancestral karyotype of most extant mosses consisted of seven chromosomes^12^, which we refer to as Ancestral Elements A-G (Fig. 1), and suggests major parts of the gene content of moss chromosomes are stable over hundreds of millions of years, similar to the “Muller Elements’’ in *Drosophila*^*13*^. Curiously, we could not detect the homeologs of the *C. purpureus* chromosomes 5 and 9 using synteny, an observation we return to below.

**Fig. 1.**
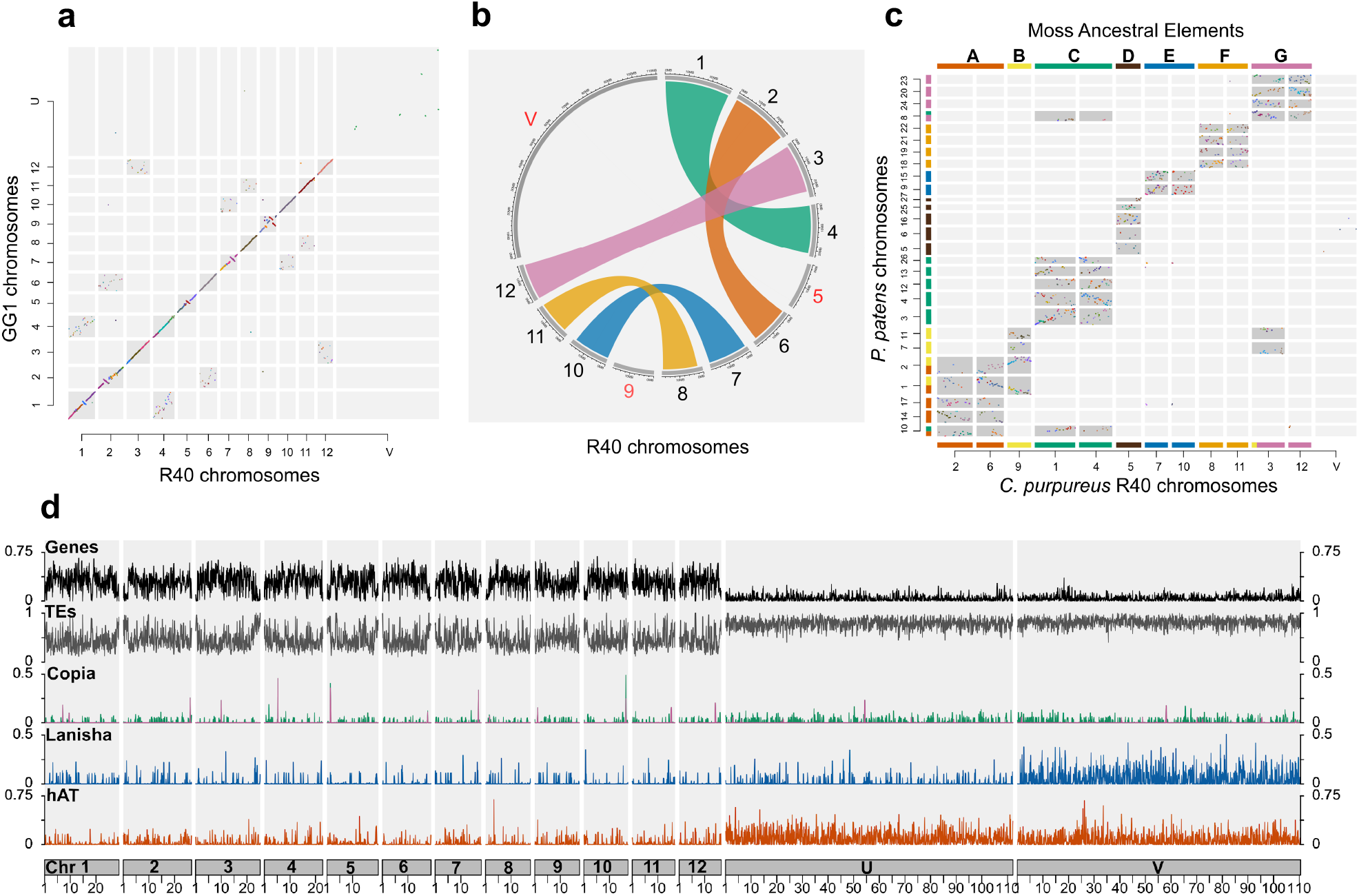
Chromosome architecture in *C. purpureus*. **a**, Dot plot of syntenic orthogroup blastp hits between *C. purpureu*s GG1 and R40 isolates, showing structural variation on autosomes and a lack of synteny across the sex chromosomes. **b**, Self-synteny plot of *C. purpureus* R40 isolate showing homeologous chromosomes from a whole-genome duplication. **c**, Dot plot of syntenic orthogroup blastp hits between *C. purpureus* R40 and *P. patens*, highlighting the seven ancestral chromosomes that we refer to as the moss ancestral elements A-G. **d**, Density plots across *C. purpureus* chromosomes (in Mb). Densities show the proportion of a 100 Kilobase (Kb) window (90 Kb jump) of each feature. Local density peaks of RLC5 Copia elements (purple Copia peaks) on each chromosome represent candidate centromeric regions, similar to *P. patens*^*12*^.

The major exception to the long-term genomic stability observed in *C. purpureus* was the sex chromosomes, which also share no obviously syntenic regions with each other or the autosomes (Fig. 1). The sex chromosomes are ∼30% of each genome (110.5 Mb on the R40 V, 112.2 Mb on the GG1 U; Fig. 1), four times the size of the largest autosome. The size is largely attributable to an increase in transposable elements (TEs), which comprise 78.2% and 81.9% of the U and V, respectively, similar to the non-recombining Y or W sex chromosomes in other systems^14^, but far more than the *C. purpureus* autosomes (mean (μ): 46.4%; *Mann-Whitney U with Benjamini and Hochberg correction* (*MWU*), autosomes to U or V *P*<2×10^−16^; Fig. 1). While some TEs have a homogeneous distribution across all chromosomes (e.g., Copia; μ: autosomes=0.8%, U=1.3%, V=1.2%; *MWU*, all pairwise comparisons *P*>0.09), the U and V chromosomes are enriched for very different classes of repeats compared to each other and the autosomes. For example, the U was enriched for hAT (μ: autosomes=2.3%, U=10.1%, V=7.4%; *MWU*, all pairwise comparisons *P*<1.5×10^−14^) and the V was enriched in a previously undescribed superfamily of cut-and-paste DNA transposons, which we refer to as Lanisha elements (μ: autosomes=1%, U=1.2%, V=5.8%; *MWU*, all pairwise comparisons *P*<1×10^−4^; Fig. 1; Supplementary Fig. 2). The distribution of repeats in *C. purpureus* and the physical proximity of the autosomes inferred from the Hi-C contact map (Extended Data Fig. 1) together highlight the enigmatic isolation of the sex chromosomes in the nucleus^15^.

Unlike other non-recombining sex chromosomes, neither the U nor V show signs of major degeneration beyond the increased TE density. Sex-linked genes used on average one more codon than autosomes (ENC), less frequently use optimal codons (fop), have a loss of preferred GC bias in the third synonymous codon position (GC3s), and have a higher rate of protein evolution (dN/dS), all consistent with weaker selection (Fig. 2; *MWU*, autosomes to U or V *P*<6×10^−6^ for all metrics). Although, notably, the U and V-linked genes were not significantly different from one another (*MWU*, ENC *P*=0.8; fop *P*=0.22; GC3s *P*=0.18; dN/dS *P*=0.73), suggesting transmission through one sex or the other has no detectable effect on purifying selection. Consistent with this observation, the U and the V possess 3,450 and 3,411 transcripts, respectively, representing ∼12% of the *C. purpureus* gene content. This stands in stark contrast to the non-recombining mammalian Y chromosome, or even other UV systems, which typically contain an order of magnitude fewer genes, at most^16–18^. These observations indicate that although suppressed recombination decreases the efficacy of natural selection, alone it is insufficient to drive gene loss on non-recombining sex chromosomes^19^.

**Fig. 2.**
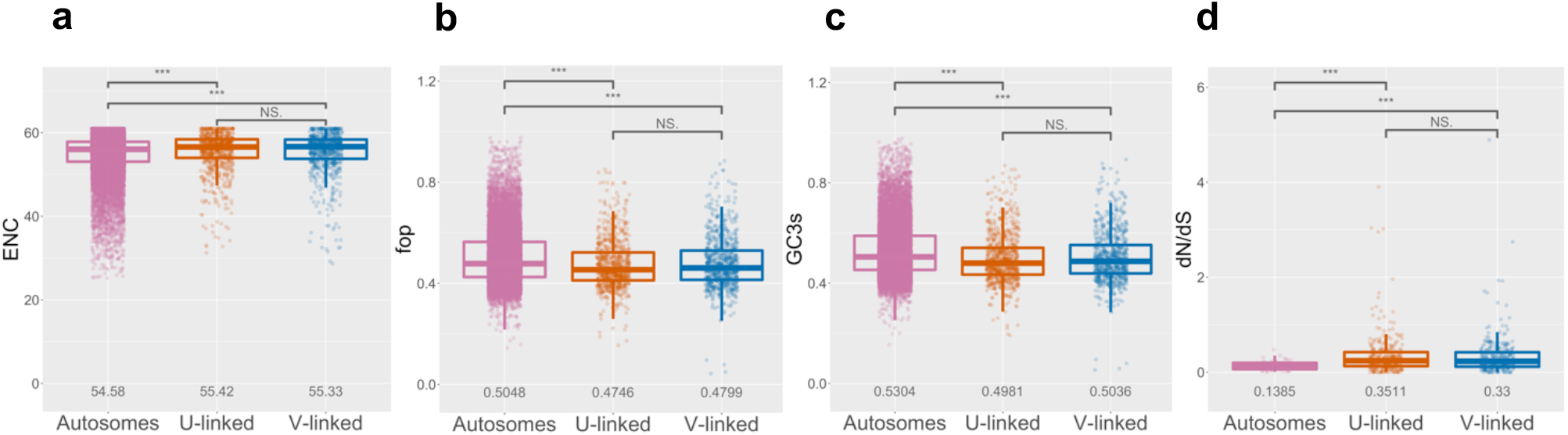
Molecular evolution of autosomal and sex-linked genes in *C. purpureus*. **a**, Autosomal genes are significantly different from U or V-linked genes in the effective number of codons (ENC) **b**, Frequency of optimal codons (fop) **c**, GC content of the third, synonymous codon (GC3s) and **d**, Protein evolution (dN/dS) (*MWU*, autosomes to U or V *P*<6×10^−6^ for all metrics, indicated by ***; numbers show means). However, U and V-linked genes were not significantly different (*MWU*, ENC *P*=0.8; fop *P*=0.22; GC3s *P*=0.18; dN/dS *P*=0.73, indicated by NS), suggesting weak but not significantly different degeneration on the U and V.

The lack of degeneration means that thousands of genes can be used to reconstruct a detailed history of gene gain on the *C. purpureus* UV sex chromosomes. Critically, the times to the most-recent common ancestor between orthologous genes on the U and V chromosomes allows us to estimate a minimum age for the sex chromosome system. To accomplish this, we used a phylogenomic approach with stringent inclusion criteria. We built 744 gene trees, 402 with U and V-linked homologs. We found most genes became sex-linked in the *C. purpureus* lineage, after the divergence from *Syntrichia princeps* (μ *Ks*=0.16; Fig. 3; Supplementary Table 7). However, 13 U-V orthologous pairs diverged at the base of the Dicranidae (μ *Ks*=0.85), and three pairs diverged prior to the split between the two diverse clades Bryidae and Dicranidae (μ *Ks*=1.64). The most ancient U-V divergence (a Zinc finger Ran binding protein of unknown function) was prior to the split between *Buxbaumia aphylla* and the remaining Bryopsida, ∼300 MYA (based on previous fossil-calibrated, relaxed-clock analyses^20^; *Ks*=2.8; Extended Data Fig. 2), making the moss sex chromosomes among the oldest known to date across Eukarya.

**Fig. 3.**
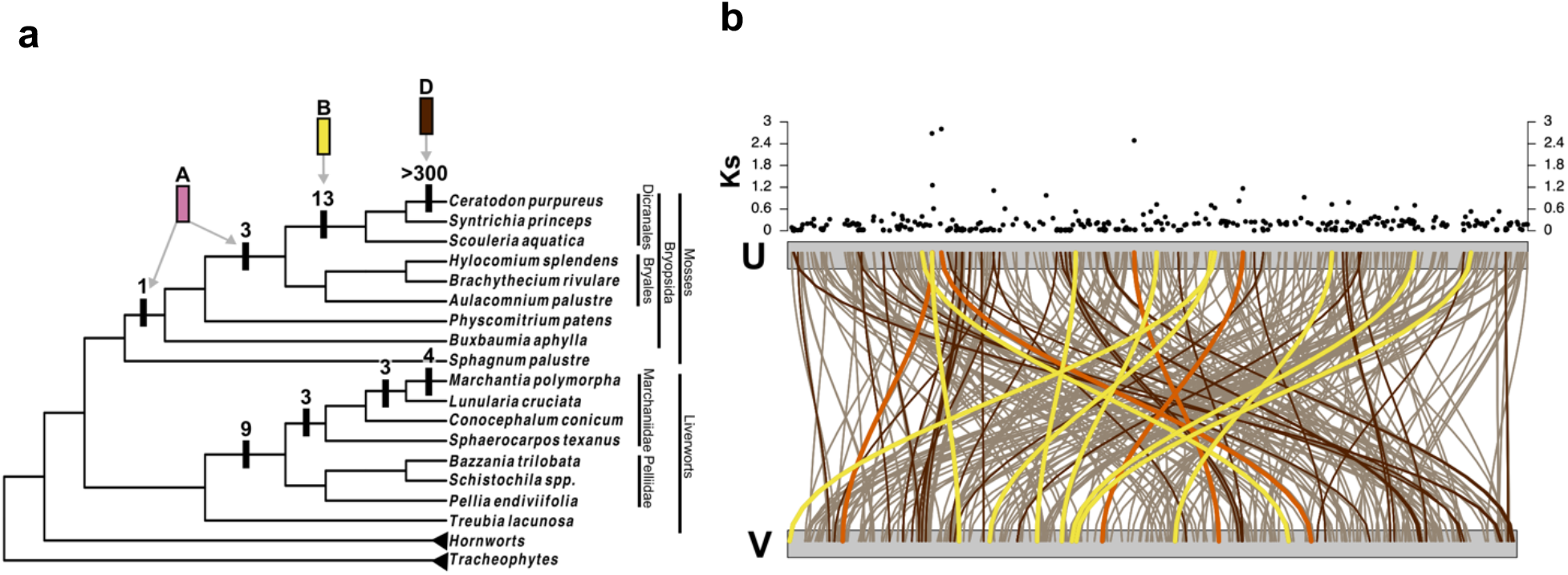
Evolutionary history of moss and liverwort sex chromosomes. **a**, Capture events of genes on moss and liverwort sex chromosomes. Numbers indicate how many extant genes were captured at the indicated branch based on the topology of the tree. The capture events in mosses can be traced back to three ancestral elements (A, B, and D), where the oldest sex-linked genes were from ancestral element A and homeologous chromosomes from B and D fused to the sex chromosomes. **b**, *Ks* of one-to-one U-V orthologs plotted on U and V sex chromosomes of *C. purpureus*. Lines connect the U-V orthologs, where colors correspond to the ancestral elements in **a**. The darker brown lines indicate genes with *Ks*≤0.02, presumably representing the most recently captured genes, which highlights the rapid rearrangement of genes on the sex chromosomes. These data, in addition to synteny (Fig. 1), also suggest a lack of a pseudoautosomal region between the *C. purpureus* U and V.

A classic signature of gene capture on sex chromosomes is the presence of strata, where neighboring genes added in the same recombination suppression event have a similar *Ks*^*21*^. However, on the *C. purpureus* sex chromosomes we found *Ks* was not associated with gene order (Fig. 3). Even genes with very low *Ks*, presumably from the most-recent recombination suppression event, were found across the entirety of the U or V, meaning gene-order was shuffled soon after the evolution of sex-linkage. To understand the mechanism by which the region of suppressed recombination acquires new genes, we combined inferences from phylogenomic analyses with the physical position of orthologs among the ancestral karyotypic elements. When we examined gene trees for the two most recent capture events, we found most homologs are from Ancestral Elements D and B (Supplementary Table 7) indicating the missing homeologous chromosomes to *C. purpureus* 5 and 9, respectively, had fused to the sex chromosomes (Fig. 3), but the scrambling of gene order had rendered them undetectable using synteny alone.

To extend this ancestral reconstruction to liverwort sex chromosomes, we examined gene trees of sex-linked genes in *M. polymorpha*^*17,22*^. Like in *C. purpureus*, we found no evidence of syntenic strata when we compared *Ks* between the U and V-linked orthologs (Supplementary Table 8). We also found evidence of four liverwort-specific capture events, with the oldest diverging ∼400 MYA, prior to the split of Marchantiidae and Pelliidae^20^ (Fig. 3; Extended Data Fig. 2). Most sex-linked genes in *M. polymorpha* (Supplementary Table 8), like two of the oldest genes in *C. purpureus* (Supplementary Table 7) have homologs from moss Ancestral Element A, leading to the remarkable suggestion that this element played a key role in sex determination early in the history of both lineages, ∼500 MYA.

A key factor explaining the retention of transcripts on non-recombining sex chromosomes is broad expression^23,24^, which in plants include the haploid phase. In transcriptomic data from multiple tissues we found more than 1,700 U and V-linked genes expressed (mean count ≥1) (Supplementary Tables 9-11), including essential components of the cytoskeleton (e.g., Tubulin) and DNA repair complexes (e.g., *RAD51*). In fact, based on an analysis of sex-biased gene expression, we found far more sex-specific sex-linked genes (i.e., only on the U or V) than significant autosomal genes (mean count ≥1, fold change ≥2, adjusted *P*≤0.05), suggesting sex-linked loci contribute more to expression differences between the sexes than do autosomes (Extended Data Table 1). Furthermore, in contrast to data from gene-poor sex chromosome systems, we found nearly all the genes in the male and female co-expression modules, including the hubs, are sex-linked (Extended Data Fig. 3; Supplementary Table 12).

The sex-specific gene expression networks are enriched for proteins with known reproductive functions across green plant lineages. For example, the male GO and KEGG terms are enriched for microtubule-based processes, which play a role in sperm production in other systems^25,26^ (Extended Data Fig. 3; Supplementary Tables 13-14). We also found both male and female co-expression modules are enriched for genes involved in circadian rhythm, like phytochrome, which are involved in flower development in *Arabidopsis thaliana*^*27*^. The male co-expression module also contained a V-specific *ABC1* gene orthologous to a V-linked copy in *M. polymorpha* (Extended Data Fig. 4), and genes in this family are involved with pollen development in angiosperms^28^. The female co-expression module contains a U-specific *RWP-RK* TF orthologous to *M. polymorpha MpRKD*, which is a component of the egg development pathway in *M. polymorpha* and *A. thaliana* and mating-type loci in green algae^16,29,30^ (Extended Data Fig. 4). The cis-acting sexual dimorphism switch *MpFGMYB*^*31*^, which promotes female development in *M. polymorpha* has orthologous U and V-linked copies in *C. purpureus* (Extended Data Fig. 4). Additionally, several other TFs or transcriptional regulators (TR) are found in the sex-specific co-expression modules (e.g., V-linked *R2R3-MYB*) or are only found on the U or V (e.g., *HD DDT, Med7*, and *SOH1*; Supplementary Table 15), together suggesting candidate regulators of sex-specific developmental processes are enriched on the *C. purpureus* UV sex chromosomes.

## Conclusions

Our analyses challenge the idea that suppressed recombination and sex-limited inheritance are sufficient to drive sex chromosome degeneration. Clearly the lack of meiotic recombination both weakens purifying selection, resulting in decreased codon bias and increased protein evolution, and facilitates massive structural variation and highly-differentiated TE accumulation between the U and V. Like in other plants, haploid gene expression in *C. purpureus* apparently slows sex chromosome degeneration, even over millions of years of suppressed recombination^23^. However, unlike flowering plants, where hermaphroditism is the norm^32^, the antiquity of dioecy in bryophytes more closely mirrors the sexual systems in animals^33,34^. Thus, the gene rich *C. purpureus* sex chromosomes provide a powerful comparative tool for studying the long-term evolution of sex-limited gene regulatory networks that govern sexual differentiation.

## Acknowledgements

The authors thank Ralph Quatrano, David Cove, and the Quatrano laboratory at Washington University in St. Louis for incubating the *C. purpureus* genome project; Bernie Hauser, Thomas Colquhoun, and Drake Garner for assisting with the hormone and light perturbations; Bernard Goffinet, Michelle Mack, Sharon Robinson, Todd Rosenstiel, Blanka Shaw, and the late Norton Miller for providing field collections. Susanne Renner, Sally Otto, Mark Kirkpatrick, and Matthew Hahn provided valuable feedback on a draft of the manuscript. The One Thousand Plant Transcriptome Initiative generously provided early access to moss and liverwort data, and the University of Florida Interdisciplinary Center for Biotechnology Research and HiPerGator provided vital technical support throughout the project. This work was supported by NSF DEB-1541005 and start-up funds from UF to SFM; microMORPH Cross-Disciplinary Training Grant, Sigma-Xi Grant-In-Aid of Research, and Society for the Study of Evolution Rosemary Grant Award to SBC; NSF DEB-1239992 to NJW; Emil Aaltonen Foundation and the University of Turku to SO; NSF DEB-1541506 to JGB and SFM. The work conducted by the U.S. Department of Energy Joint Genome Institute was supported by the Office of Science of the U.S. Department of Energy under Contract No. DE-AC02-05CH11231.

## Author contributions

Conceptualization: SFM; Data curation: SBC, KB; Formal analysis: SBC, JJ, SS, JTL, FM, AS, GPT, NFP, SAR; Funding acquisition: SBC, SH, NJW, JGB, SAR, SFM; Investigation: SBC, JJ, ACP, SS, JTL, FM, AS, GPT, NFP, CC, MW, AL, CD, CS, JCM, REC, LMK, SO, SH, JBL, JGB, SAR, SFM; Methodology: SBC, ACP, JJ, JS, SFM; Project administration: SBC, KB, JG, JS, SFM; Resources: NJW, MJG, JG, JS, SFM; Supervision: MGJ, SAR, JG, JS, SFM; Visualization: SBC, JJ, JTL, AS, GPT; Writing – original draft: SBC, SFM; Writing – review and editing: SBC, ACP, JJ, JTL, FM, AS, GPT, CAS, LMK, JGB NJW, MGJ, SAR, JG, JS, SFM. All authors approve of the final draft of this manuscript.

## Competing interests

The authors declare no competing interests.

## Data availability

DNA and RNA data for this manuscript can be found under NCBI BioProjects listed in Supplementary Table 5. The R40 and GG1 v1.0 genome assemblies can be found on NCBI GenBank under the accessions JACMSA000000000 and JACMSB000000000, respectively. The genome assemblies and annotations can also be found on Phytozome (https://phytozome-next.jgi.doe.gov/). The Lanisha transposable element has been deposited in NCBI GenBank under MT647524. The published sequence data used in this study can be found in Supplementary Table 16. Supporting documents for the phylogenomic analysis of sex-linked genes can be found on Dryad under https://doi.org/10.5061/dryad.v41ns1rsm.

## Methods

### Isolate collection and tissue culture

All *C. purpureus* tissue used in this study was isolated from a single-spore^35^, from field-collected sporophytes (Supplementary Table 5)^10,36^. In-depth methods for tissue generation for DNA and RNA, library preparation, and sequencing can be found in the Supplementary Methods.

### Genome assemblies

We sequenced *C. purpureus* (var. GG1 and var. R40) using a whole-genome shotgun sequencing strategy and standard sequencing protocols. Sequencing reads were collected using Illumina, PacBio, and Sanger platforms. Illumina, PacBio, and Sanger reads were sequenced at the Department of Energy (DOE) Joint Genome Institute (JGI) in Walnut Creek, California and the HudsonAlpha Institute in Huntsville, Alabama. Illumina reads were sequenced using the Illumina HiSeq-2000 and X10 platform and the PacBio reads were sequenced using the SEQUEL I platform. Sanger BACs were sequenced using an ABI 3730XL capillary sequencer. For both GG1 and R40, one 400bp insert 2×150 Illumina fragment library (133.14x for GG1, 146.45x for R40) was sequenced along with one 2×150 Dovetail HiC library (252.86x GG1, 442.71x R40) (Supplementary Table 1). Prior to assembly, Illumina fragment reads were screened for phix contamination. Reads composed of >95% simple sequence were removed. Illumina reads <50bp after trimming for adapter and quality (q<20) were removed. For the PacBio sequencing, a total of 8 PB chemistry 2.1 cells (10 hour movie time) were sequenced each for GG1 and R40 on Sequel 1 with a raw sequence yield of 39.82 Gb (GG1) and 46.24 Gb (R40) with a total coverage of 113.77x (GG1) and 132.11x (R40) (Supplementary Table 2). Finally, a total of 1,032 BAC clones sequenced with Illumina indexed libraries were used for patching the final chromosome gaps.

#### Genome assembly and construction of pseudomolecule chromosomes

Improved versions 1.0 of the *C. purpureus* (var. GG1 and var. R40) assemblies were generated by assembling the 4,195,510 PacBio GG1 reads (113.77x sequence coverage) and 5,238,148 PacBio reads R40 (132.11x sequence coverage) separately using the MECAT assembler^37^ and subsequently polished using QUIVER^38^. For GG1, this produced 637 scaffolds (637 contigs), with a contig N50 of 1.2 Mb, 475 scaffolds larger than 100 Kb, and a total genome size of 347.1 Mb (Supplementary Table 3). For R40, this produced 731 scaffolds (731 contigs), with a contig N50 of 1.1 Mb, 497 scaffolds larger than 100 Kb, and a total genome size of 361.3 Mb (Supplementary Table 3).

Hi-C scaffolding using the JUICER pipeline^39^ was used to identify misjoins in the initial MECAT assembly. Misjoins were characterized as a discontinuity in the GG1 or R40 linkage group. A total of 73 misjoins were identified and resolved in GG1 and 64 in R40. The resulting broken contigs were then oriented, ordered, and joined together into 13 chromosomes (12 autosomal and 1 sex chromosome designated as “U” in the GG1 release and 12 autosomal and 1 sex chromosome designated as “V” in the R40 release) using both the map, as well as the Hi-C data. A total of 579 joins were made in GG1 and 625 in R40 during this process. Each chromosome join is padded with 10,000 Ns. Significant telomeric sequence was identified using the (TTTAGGG)_n_ repeat, and care was taken to make sure that it was properly oriented in the production assembly. The remaining scaffolds were screened against bacterial proteins, organelle sequences, GenBank nr and removed if found to be a contaminant. For GG1, a set of 1,032 BAC clones (107.8 Mb total sequence) sequenced with Illumina indexed libraries were used to patch remaining gaps in the chromosomes. Clones were aligned to the chromosomes using BLAT^40^, and clone contigs crossing gaps were used to form patches. A total of 35 gaps were patched.

Finally, homozygous SNPs and INDELs were corrected in the release consensus sequence using ∼88x of Illumina reads (2×150, 400bp insert) by aligning the reads using bwa mem^41^ and identifying homozygous SNPs and INDELs with the GATK’s UnifiedGenotyper tool^42^. A total of 108 homozygous SNPs and 5,291 homozygous INDELs in GG1 and 19 homozygous SNPs and 867 homozygous INDELs in R40 were corrected in the release. The final version 1.0 GG1 release contains 349.5 Mb of sequence (1.3% gap), consisting of 558 contigs with a contig N50 of 1.4 Mb and a total of 97.9% of assembled bases in chromosomes. The final version 1.0 R40 release contains 358.0 Mb of sequence (1.2% gap), consisting of 601 contigs with a contig N50 of 1.4 Mb and a total of 98.3% of assembled bases in chromosomes.

Completeness of the euchromatic portion of the version 1.0 GG1 and 1.0 R40 assemblies was assessed by aligning an rnaSEQ library (library code GNGZB for GG1, GNGZC for R40). The aim of this analysis is to obtain a measure of completeness of the assembly, rather than a comprehensive examination of gene space. The transcripts were aligned to the assembly using GSNAP^43^. The alignments indicate that 96.88% of the GG1 RNAseq reads aligned to the version 1.0 GG1 release and 97.01% of the R40 RNAseq reads aligned to the version 1.0 R40 release.

#### Construction of the scaffold assembly

A total of 4,195,510 PacBio reads (113.77x) in GG1 and 5,238,148 PacBio reads (132.11x) in R40 were assembled using MECAT^37^, and formed the starting point of the version 1.0 release for each. The 310,662,272 Illumina sequence reads (133.14x sequence coverage) in GG1 and 353,932,084 Illumina sequence reads (146.45x sequence coverage) in R40 were used for fixing homozygous snp/indel errors in the consensus. A total of 310,662,272 Hi-C reads (252.86x sequence coverage) in GG1 and 1,062,837,932 Hi-C reads (442.71x sequence coverage) in R40 were used for chromosome construction.

#### Screening and final assembly release

Scaffolds that were not anchored in a chromosome were classified into bins depending on sequence content. Contamination was identified using blastn against the NCBI nucleotide collection (NR/NT) and blastx using a set of known microbial proteins. In GG1, additional scaffolds were classified as repetitive (>95% masked with 24mers that occur more than 4 times in the genome) (16 scaffolds, 482.8 Kb), chloroplast (1 scaffold, 158.7 Kb), and low quality (>50% unpolished bases post polishing, 3 scaffolds, 48.3 Kb) In R40, additional scaffolds were classified as repetitive (>95% masked with 24mers that occur more than 4 times in the genome) (12 scaffolds, 489.6 Kb), chloroplast (1 scaffold, 50.2 Kb), and low quality (>50% unpolished bases post polishing, 6 scaffolds, 236.8 Kb). Resulting final statistics are shown in Supplementary Table 4.

#### GG1 assessment of assembly accuracy

A set of 17 finished contiguous Sanger BAC clones >100Kb were selected in order to assess the accuracy of the assembly. A range of variants were detected in the comparison of the BAC clones and the assembly. In 14 of the BAC clones, the alignments were of high quality (< 0.05% bp error) with an example being given in Supplementary Fig. 1. All dot plots were generated using Gepard^44^. The remaining 3 BACs indicate a higher error rate due mainly to their placement in more regions containing tandem repeats (Supplementary Fig. 2). The overall bp error rate in the BAC clones is 0.016% (269 discrepant bp out of 1,599,605 bp).

### Genome Annotations

Transcript assemblies were made from ∼1.5 billion pairs of 2×150 stranded paired-end Illumina RNA-seq reads from *C. purpureus* GG1 and ∼1.6 billion pairs from *C. purpureus* R40 using PERTRAN (Shu, unpublished), PERTRAN assemblies from G100m_X_G150f_Sporo reads on the *C. purpureus* GG1 or R40 genome, and filtered open reading frames (ORFs) from Trinity assemblies from stranded paired-end Illumina reads from additional *C. purpureus* cultivars (Chile, Dur, Ecud, Ren, and UCONN; Supplementary Table 5) 180,954 (GG1) and 194,414 (R40) transcript assemblies were constructed using PASA^45^ from RNA-seq transcript assemblies above and a bit of *C. purpureus* ESTs. Loci were determined by transcript assembly alignments and/or EXONERATE alignments of proteins from *Arabidopsis thaliana*^*46*^, soybean^47^, *Setaria viridis*^*48*^, grape^49^, *Sphagnum magellanicum, Physcomitrium patens*^*12*^, *Selaginella moellendorffii*^*50*^, *Chlamydomonas reinhardtii*^*51*^, filtered Trinity assembly ORFs described above, high-confidence gene models from the first round of *C. purpureus* R40 gene call, uniprot Bryopsida and Swiss-Prot proteomes to repeat-soft-masked *C. purpureus* GG1 genome using RepeatMasker^52^ with up to 2K BP extension on both ends unless extending into another locus on the same strand. Repeat library consists of De Novo repeats by RepeatModeler ^53^ on *C. purpureus* GG1 genome and repeats in RepBase. Gene models were predicted by homology-based predictors, FGENESH+^54^, FGENESH_EST (similar to FGENESH+, EST as splice site and intron input instead of protein/translated ORF), and EXONERATE^55^, PASA assembly ORFs (in-house homology constrained ORF finder) and from AUGUSTUS via BRAKER1^56^. The best scored predictions for each locus are selected using multiple positive factors including EST and protein support, and one negative factor: overlap with repeats. The selected gene predictions were improved by PASA. Improvement includes adding UTRs, splicing correction, and adding alternative transcripts. PASA-improved gene model proteins were subject to protein homology analysis to above mentioned proteomes to obtain Cscore and protein coverage. Cscore is a protein BLASTP score ratio to MBH (mutual best hit) BLASTP score and protein coverage is the highest percentage of protein aligned to the best of homologs. PASA-improved transcripts were selected based on Cscore, protein coverage, EST coverage, and its CDS overlapping with repeats. The transcripts were selected if its Cscore is larger than or equal to 0.5 and protein coverage larger than or equal to 0.5, or it has EST coverage, but it’s CDS overlapping with repeats is less than 20%. For gene models whose CDS overlaps with repeats for more than 20%, its Cscore must be at least 0.9 and homology coverage at least 70% to be selected. The selected gene models were subject to Pfam analysis and gene models whose protein is more than 30% in Pfam TE domains were removed and weak gene models. Incomplete gene models, low homology supported without fully transcriptome supported gene models and short single exon (<300 bp CDS) without protein domain nor good expression gene models were manually filtered out.

### Synteny analysis within *C. purpureus* and between *P. patens*

We ran the default GENESPACE pipeline^57^ with a minimum block size of 5 genes and a maximum gap / search radius of 15 genes. In short, GENESPACE runs orthofinder on synteny-constrained blastp hits. This offers higher stringency when exploring highly diverged genomes (or ancient whole-genome duplications) by removing high scoring, but randomly distributed, blast hits.

### Ks-plot analysis to identify the *C. purpureus* whole-genome duplication

Whole-genome duplications (WGD) were detected with conventional *Ks* plot analyses. We used the wgd pipeline^58^. An all-by-all BLASTP search^59^ was performed for the *C. purpureus* GG1 and R40 genomes as well as *P. patens* and *M. polymorpha*. Paralogs were clustered with MCL^60^. For each cluster, all pairwise *Ks* estimates were obtained from PAM^61^ with the GY94 model with F3×4 equilibrium codon frequencies^62^. Hierarchical clustering was used to reduce redundant comparisons and obtain node-averaged *Ks* estimates. This process was repeated for syntenic paralogs too, which were obtained from I-ADHoRe v3.0 with default settings^63^ based on all-by-all BLASTP results. Orthologous gene divergences used reciprocal best BLASTP hits between *C. purpureus* and *P. patens*.

Peaks in *Ks* plots can be identified visually, but we also applied mixture models that were selected by the difference in BIC scores, such that a difference less than 3.2 is used as a stopping criterion. Mixture models were implemented with the *bic*.*test*.*wgd* function available on GitHub (https://github.com/gtiley/Ks_plots). Mixture models can be problematic in their interpretation due to over-fitting, therefore we looked for peaks that were consistently detected across models and the maximum *Ks* value allowed^64^. When analyzing all paralogs, a single prominent peak was observable in *C. purpureus* with a mean between a *Ks* of 0.65 and 0.97 in GG1 and a *Ks* between 0.68 and 0.74 in R40 (Supplementary Table 6). The more consistent results in R40 imply more paralogs from this WGD event have survived on the V chromosome compared to the U chromosome. This WGD post-dates the divergence of *C. purpureus* and *P. patens* (Fig. S12). This is determined by visual inspection but agrees with previous analyses of WGD in both *C. purpureus* and *P. patens*^*10–12*^. The presence of a single WGD that occurred in *C. purpureus* following divergence from *P. patens* is supported by analyses of syntenic paralogs as well (Extended Fig. 1), which suggest slightly more recent WGD ages (Supplementary Table 6). However, analyses of syntenic paralogs from *P. patens* supported the presence of two WGDs following divergence from *C. purpureus* (Supplementary Table 6), similar to previous findings when using syntenic data^12^ compared to all paralogs from genomic or transcriptomic data^11,65^.

*Ks-*plot analyses are provocative of older WGD events that pre-date the divergence of *C. purpureus* and *P. patens*. Notably low numbers of syntenic paralogs are evident between *Ks* of 3.0 and 4.0; although, the same is true for *M. polymorpha* that putatively has no history of ancient WGD. Any identifiable peaks in *Ks* plot analyses are too speculative given lack of evidence from mixture models, and nor do their existence affect our proposed model of karyotype evolution. It should be noted though that analyses of gene trees that reconcile duplication and loss events onto a species tree have implied a shared large-scale duplication event shared by *C. purpureus* and *P. patens* (“B3”^11^) and an even older event shared by all mosses (“B2”^11^). Testing such ancient hypotheses is beyond the scope of *Ks* plot analyses, even with syntenic data. Rather, macrosyntenic evidence from more moss species, such as *Sphagnum fallax*, will be needed to identify the presence of expected syntenic ratios among genes, similar to the identifiable 1:4 ratios between *C. purpureus* and *P. patens* investigated here.

### Transposable element annotation

We combined R40 assembly (autosomes and V) with the U sex chromosome assembled from GG1 to run *de novo* repeat detection using the TEdenovo pipeline from the REPET package (v2.4)^66^. Parameters were set to consider repeats with at least 5 copies. We obtained a library of 4,699 consensus sequences that was filtered to keep only those that are found at least once as full length copy in the combined assembly and we retained 2,523 of them. This library of consensus sequences was then used as digital probe for whole genome annotation by the TEannot^67^ pipeline from the REPET package v2.4. Threshold annotation scores were determined for each consensus as the 99th percentile of the scores obtained against a randomized sequence (reversed input, not complemented and masked using Tandem Repeats Finder with parameters 2 7 7 80 10 70 10^68^). The library of consensus sequences was classified using PASTEC followed by manual curation^69^. To improve classification, remote homology detection was performed using HH-suite3^70^. For the density plot of genes and TEs (Fig. 1), we calculated the proportion of coverage of each feature in a 100 Kb window with a 90 Kb jump using Bedtools (v2.27.)^71^. These results were plotted in R (v3.5.3)^72^ using the package karyoploteR (v1.8.8)^73^ and edited in Inkscape (v0.92.2) (https://inkscape.org/en/) (ceratodon_genome_plots.R, https://doi.org/10.5061/dryad.v41ns1rsm). To examine differences in enrichment between the autosomes, U, and V we ran a pairwise Mann-Whitney U test with a Benjamini and Hochberg correction for multiple tests^74,75^ using the sliding window densities (n_Auto_=2736, n_U_=1247, n_V_=1229).

### Transcription factor and regulator annotation

Transcription associated proteins (TAPs) comprise transcription factors (TFs, acting in sequence-specific manner, typically by binding to cis-regulatory elements) and transcriptional regulators (TRs, acting on chromatin or via protein-protein interaction). We classified all *C. purpureus* proteins into 122 families and sub-families of TAPs by a domain-based rule set^76,77^. We compared this genome-wide classification with relevant organisms. All proteins in which a domain was found are listed with their family assignment. In cases when the domain composition does not allow an unambiguous assignment they are assigned no_family_found.

### Gene expression and co-expression

Gene expression and co-expression analyses were done using three male-female sibling pairs (n_isolates_=6, 3 of each sex) at gametophore and protonemal stages (n_stages_=2) in triplicate (n_replicates_=3) (Supplementary Table 5; see Supplementary Methods for details on tissue conditions). Raw reads were filtered for contaminants and adapters removed using BBDuk (v38.00) (Bushnell, http:\\bbtools.jgi.doe.gov). This included removing reads with 93% identity to human, mouse, dog, or cat or align to common microbial references. Further filtering removed reads with any ‘N’s, an average quality of 10, or a length <50 or 33% of the full read length. Adapters were trimmed and reads were right-quality-trimmed if quality was below 6. Paired-end reads were split into forward and reverse reads (novaseq_FASTQ_de_interlacer.pl, https://doi.org/10.5061/dryad.v41ns1rsm). Reads were further filtered for quality using Trimmomatic (v0.36)^78^ using leading and trailing values of 3, window size of 10, quality score of 30, and minimum length of 40. We assessed the quality of the remaining reads using fastqc (v0.11.4) (Andrews, 2010).

Filtered reads were mapped using HISAT2 (v2.1.0)^79^ to the *C. purpureus* R40 genome (autosomes and V sex chromosome) concatenated with the GG1 U sex chromosome. We hard masked the U chromosome for males, and the V for females, using Bedtools (v2.27.1)^71^ maskfasta^80^. Genes greater than 300bp were assembled using StringTie (v1.3.3)^81^, gene counts were extracted using StringTie’s prepDE.py script (http://ccb.jhu.edu/software/stringtie/index.shtml?t=manual#deseq),and gene IDs renamed (using mstrg_prep.pl, https://gist.github.com/gpertea/b83f1b32435e166afa92a2d388527f4b).

Only genes matching the original genome annotation file were used for co-expression analyses below. To identify differentially-expressed genes (DEGs), we used DESeq2 (v1.22.2^82^) where we contrasted males and females at both the protonemal and gametophore stages. For autosomal genes, we removed those with baseMean<1, a log_2_ fold change<1, and an adjusted p-value <0.05. For sex-linked genes, we calculated the mean normalized count across only males or females for protonema and gametophore separately. To identify which sex-linked genes were sex-specific we used the output from Orthofinder below.

#### Co-expression network construction and module detection

Weighted gene co-expression networks were constructed using the WGCNA R package (v1.69)^83^ with genes expression data normalized using variance stabilizing transformation from the DESeq2 R package (v1.26.0)^84^. The data retained after filtering genes showing low expression levels (minimum read count = 6 and minimum total read count = 10) were used to construct co-expression network modules using the block-wise network construction procedures. Briefly, pairwise Pearson correlations between each gene pair were weighted by raising them to power (β). To select a proper soft-thresholding power, the network topology for a range of powers was evaluated and appropriate power was chosen that ensured an approximate scale-free topology of the resulting network.

The pairwise weighted matrix was transformed into topological overlap measure (TOM). And the TOM-based dissimilarity measure (1 – TOM) was used for hierarchical clustering and initial module assignments were determined using a dynamic tree-cutting algorithm. Pearson correlations between each gene and each module eigengene, referred to as a gene’s module membership, were calculated and module eigengene distance threshold of 0.25 was used to merge highly similar modules. Top 10 hub genes in each module were identified based on module membership. These co-expression modules were assessed to determine their correlation with expression patterns distinct to conditions. Interesting modules having significant relationships with conditions, such as sex, were visualized using the igraph (v1.2.5)^85^ and ggnetwork (v0.5.8)^86^ R packages and in order to focus on the relevant gene pair relationships, network depictions were limited to an adjacency threshold of 0.2 and the top 3000 edges/interactions between nodes/gene models.

#### GO and KEGG pathway enrichment analysis

Gene Ontology (GO) enrichment analysis was carried out using topGO, an R Bioconductor package (v2.38.1)^87^ with Fisher’s exact test; only GO terms with a P <0.05 were considered significant. To identify redundant GO terms, semantic similarity among GO terms were measured using Wang’s method implemented in the GOSemSim, an R package (v2.12.1)^88^. KEGG ^89^ pathway enrichment analysis was performed based on hypergeometric distribution test and pathways with P <0.05 were considered enriched.

### Phylogenomic analyses of moss and liverwort sex chromosomes

The genome and transcriptome lines used for phylogenomic analyses can be found in Supplementary Table 16^11,12,17,50,90–93^. For all RNA seq data, we filtered for quality using Trimmomatic (v0.36)^78^ using leading and trailing values of 3, a window size of 10, a quality score of 30, and a minimum length of 40. We assessed the quality of the remaining reads using fastqc (v0.11.4) (Andrews, 2010). To *de novo* assemble genes, we used Trinity (vr20170205-2.4.0)^94^ following default parameters (the exception being with C. *purpureus*, for which used –SS_lib_type RF). We next determined the single best open reading frame using TransDecoder (v5.0.2)^95^. Our reading frames were checked first against pFam (v32.0)^96^ and if no hit was found the frame was determined by Transdecoder. To reduce protein redundancy, we next ran our open reading frames through CD-HIT (4.6.3)^97,98^ using a 0.99 threshold.

We first found orthogroups for the in-frame genes using Orthofinder (v2.2.0)^99,100^. We built gene trees for genes annotated on the *M. polymorpha* and *C. purpureus* sex chromosomes by first filtering clusters for at least eight species present in the tree (orthogroup_filter.pl, https://doi.org/10.5061/dryad.v41ns1rsm). For these clusters, we wrote FASTA files for both amino acid and cds files of genes clustered within an orthogroup (fasta_from_OrthoFinder.pl, https://doi.org/10.5061/dryad.v41ns1rsm). We next aligned our amino acid fasta files using MAFFT (v7.407)^101^. We back translated our alignments to DNA using pal2nal (v14)^102^.

Alignments were filtered for column occupancy of 0.5 using trimal (v1.2)^103^ and filtered to remove any sequences less than 300bp (alignment_length_filter.pl, https://doi.org/10.5061/dryad.v41ns1rsm). These final alignments were used to build bootstrapped trees using RAxML (v8.2.8)^104^ using the GTRGAMMA model and 100 bootstrap replicates. We visually analyzed trees to determine when genes became sex-linked. To accomplish this, we identified the clades which contained annotated U and V-linked genes and determined the most-distantly related species found in the same clade (e.g., Extended Data Fig. 2). All trees and alignments can be found on Dryad under https://doi.org/10.5061/dryad.v41ns1rsm. All tree plots were made using ggtree (v1.14.6)^105,106^ in R (v3.5.3)^72^ and edited in Inkscape (v0.92.2) (https://inkscape.org/en/) (ceratodon_genome_plots.R, https://doi.org/10.5061/dryad.v41ns1rsm).

To identify the Ancestral Element from which sex-linked genes descended, trees with one-to-one U-V orthologs were rooted using *Azolla, Salvinia, Selaginella, Takakia*, or *Sphagnum* as an outgroup (in this order of preference) using newick utils (v1.6)^107^ and only the longest isoform within a clade for the same sample was retained (edlwtre2.pl, https://doi.org/10.5061/dryad.v41ns1rsm). To determine the closest *P. patens* gene, we used a custom python script (physco_outgroup.py, https://doi.org/10.5061/dryad.v41ns1rsm), which used ETE3^108^ to first intentify the sex-linked genes then finding the closest *P. patens* gene based on brach length. For these genes, we also determined if a paralogous *C. purpureus* chromosome 5 paralog was present and reported only those that clearly showed gene duplication, presumably from the whole-genome duplication event.

### Protein evolution

To examine protein evolution of sex-linked and autosomal genes, we first pruned the trees described above at the closest *P. patens* homolog (prune_tree.py, https://doi.org/10.5061/dryad.v41ns1rsm)^108^. For genes that had a *C. purpureus* chromosome 5 homolog, the R40 and GG1 leaves were identified instead and pruned at the closest homolog in *P. patens*. The chromosome 5 homologs were used to assess dN/dS on autosomal genes in *C. purpureus* and were specifically targeted given the recent fusion of the chromosome 5 homeolog to the sex chromosomes. All other copies of a *C. purpureus* gene were removed and which copy of a gene to keep for all other species was chosen at random. To get dN/dS ratios we used PAML (v4.9a^61^; additional scripts for this analysis in https://doi.org/10.5061/dryad.v41ns1rsm). For the sex-linked gene trees, we allowed the U and V to evolve at different rates than the rest of the tree. For the chromosome 5 homologs, the GG1 and R40 branches could evolve at a different rate than the rest of the tree. dN/dS values > 5 were removed from further analyses. To determine if there is a significant difference in dN/dS on autosomal, U, and V-linked genes we ran a pairwise Mann-Whitney U test with a Benjamini and Hochberg correction for multiple tests^74,75^ (n_auto_=61, n_U_=314, n_V_=315).

### Ka Ks analysis

FASTA files of in-frame *C. purpureus* and *M. polymorpha* sex-linked genes were aligned (see above) and converted to axt format (array_hash_extractor_fasta_unlock_ks.pl and aln_to_axt.pl, https://doi.org/10.5061/dryad.v41ns1rsm). *Ka, Ks*, and *Ka*/*Ks* were calculated using KaKs Calculator (v2.0)^109^ using the Goldman and Yang model^62^ on only one-to-one UV orthologs. *Ks* was plottred on the U and V sex chromosomes (Fig. 3) in R (v3.5.3)^72^ using karyoploteR (v1.8.8)^73^ and edited in Inkscape (v0.92.2) (https://inkscape.org/en/). One gene with *Ks* >3 but coalescence in *C. purpureus* was removed from the plot (ceratodon_genome_plots.R, https://doi.org/10.5061/dryad.v41ns1rsm).

### Codon analyses

To analyze codon-usage biases we used CodonW (v1.4.2; J. Penden https://sourceforge.net/projects/codonw). We first removed any gene that had no expression to remove potential pseudogenes (gene expression methods below). We also removed genes with less than 200 codons to reduce the variance around calculated codon values using a custom PERL script (alignment_length_filter.pl, https://doi.org/10.5061/dryad.v41ns1rsm^110^). We ran a correspondence analysis on autosomes, U, and V-linked genes together to determine the optimal codons in *C. purpureus*. We next determined the frequency of optimal codon usage (fop^111^), effective number of codons (ENC), and GC content of the third synonymous position of a codon (GC3s) on autosomes, U, and V-linked genes separately. To determine if there is a significant difference between fop, ENC, and GC3s between autosomes, U, and V we ran a pairwise Mann-Whitney U test with a Benjamini and Hochberg correction for multiple tests^74,75^ in R (v3.5.3^112^) (n_Auto_=15,677, n_U_=797, n_V_=736) and plotted the results (Fig. 2) using ggplot2 (v3.2.1^113^) using default box-plot elements (ceratodon_genome_plots.R, https://doi.org/10.5061/dryad.v41ns1rsm).

## Extended Data

**Extended Data Fig. 1.**
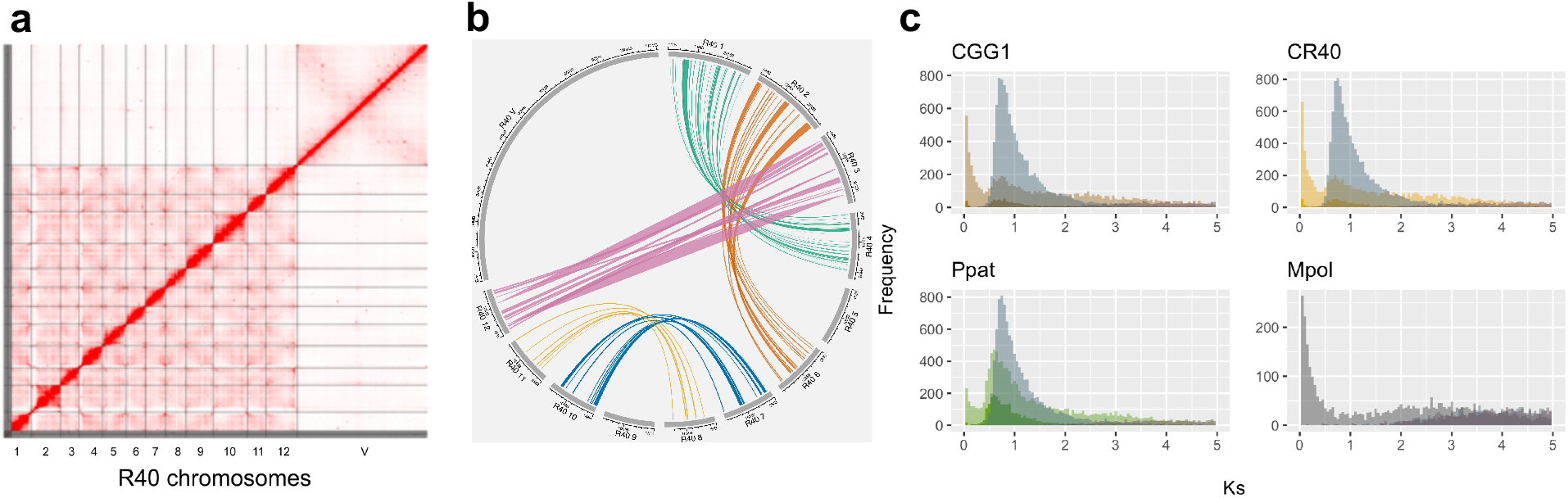
The *C. purpureus* genome assembly and whole-genome duplication analyses. **a**, Hi-C contact map for the *C. purpureus* R40 isolate, highlighting the chromosome-scale genome assembly. **b**, Collinear regions identified in R40, highlighting the homeologous chromosomes from a whole-genome duplication; synteny plots in GG1 show the same patterns (data not shown). **c**, *Ks* plots for *C. purpureus GG1* and R40, *P. patens*, and *M. polymorpha*. Node-averaged *Ks* values are shown for all paralogs (in the lighter color) as well as syntenic paralogs (the darker color). The third, blue-grey distribution are orthologs between *C. purpureus and P. patens*. The *M. polymorpha Ks* plot has no syntenic paralogs and shows *M. polymorpha* – *C. purpureus* orthologs in blue-grey as well *M. polymorpha* – *P. patens* orthologs in purple, which overlap and become indistinguishable.

**Extended Data Fig. 2.**
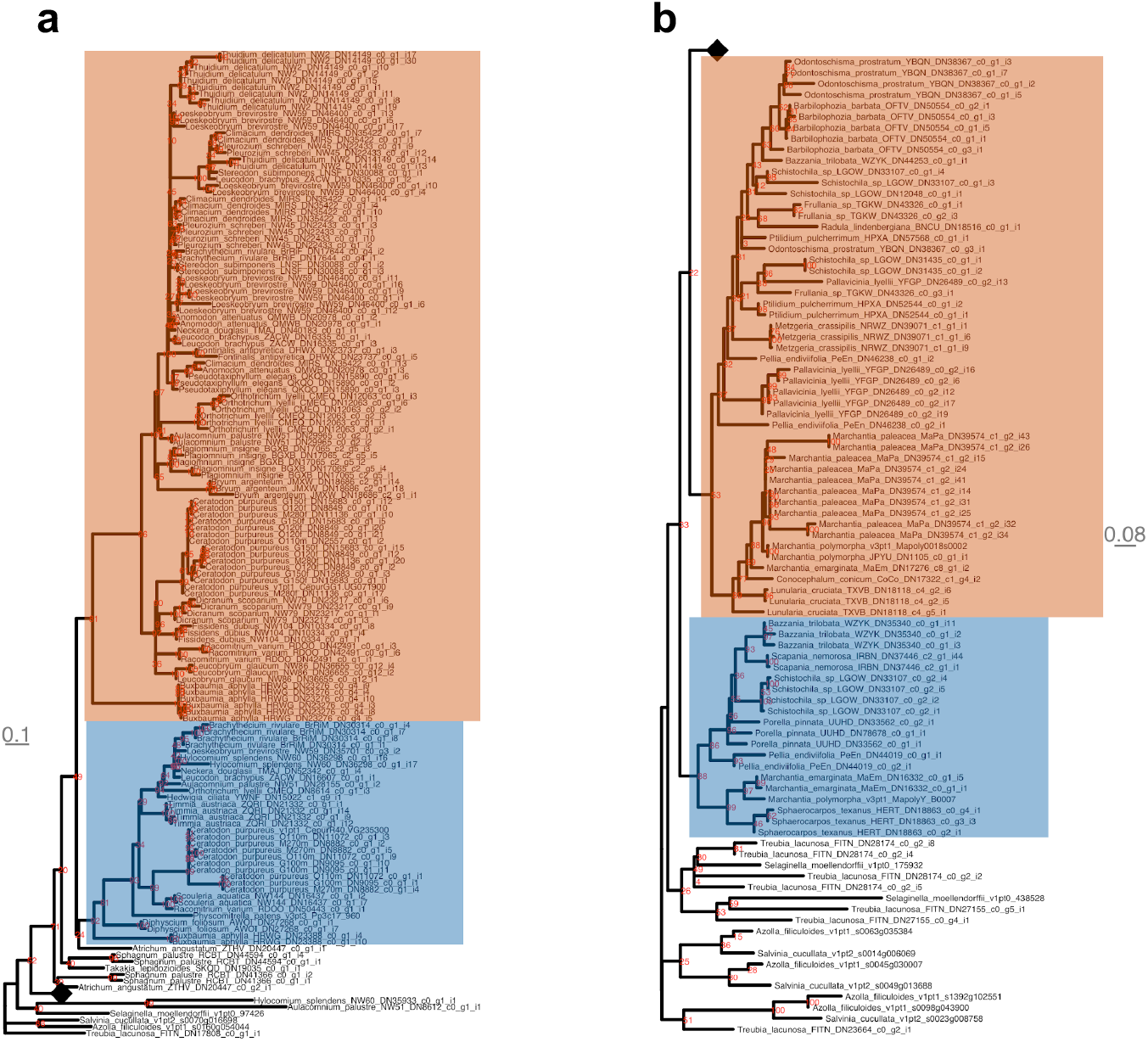
Phylogeny of oldest sex-linked gene identified in mosses and liverworts. Trees were built using Maximum-likelihood and bootstrap support is shown in red. Trees were rooted using ferns (*Azolla* or *Salvinia*) as the outgroup. Black diamonds represent collapsed clades and the branch-length scale is in gray. The clades highlighted in blue are V-linked (male), orange are U-linked (female). **a**, Ancient sex-linked gene in mosses. The tree includes sex-specific data for *C. purpureus* and distantly-related *Brachythecium rivulare*. Evidence for the ancient sex-linkage includes isolates of the same sex being more closely related than the other sex of the same species (e.g., *C. purpureus* GG1 is more closely related to BrRiF and R40 more-closely related to BrRiM). **b**, One of the oldest sex-linked genes in liverworts. Eight other trees in liverworts show a similar topology (see Supplementary Table 8).

**Extended Data Fig. 3.**
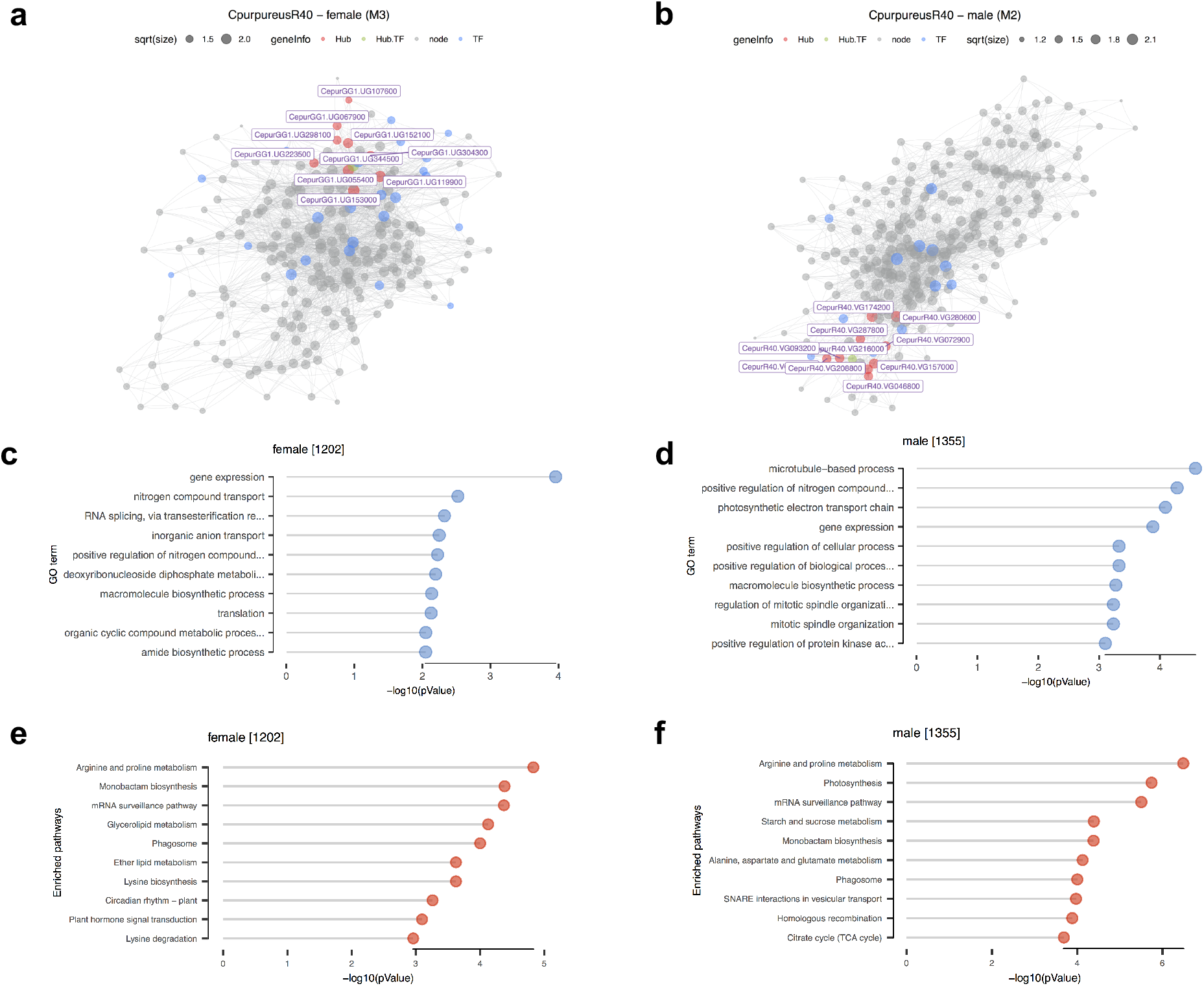
Co-expression, GO, and KEGG enrichments for female and male modules. **a**, Co-expression modules for females. **b**, Co-expression modules for males. **c**, female module GO enrichment. **d**, male module GO enrichment. **e**, female module KEGG enrichment. **f**, male module KEGG enrichment. HUBs for each module are identified in red and with the gene name shown. Transcription factors (TF) are in blue and nodes in gray.

**Extended Data Fig. 4.**
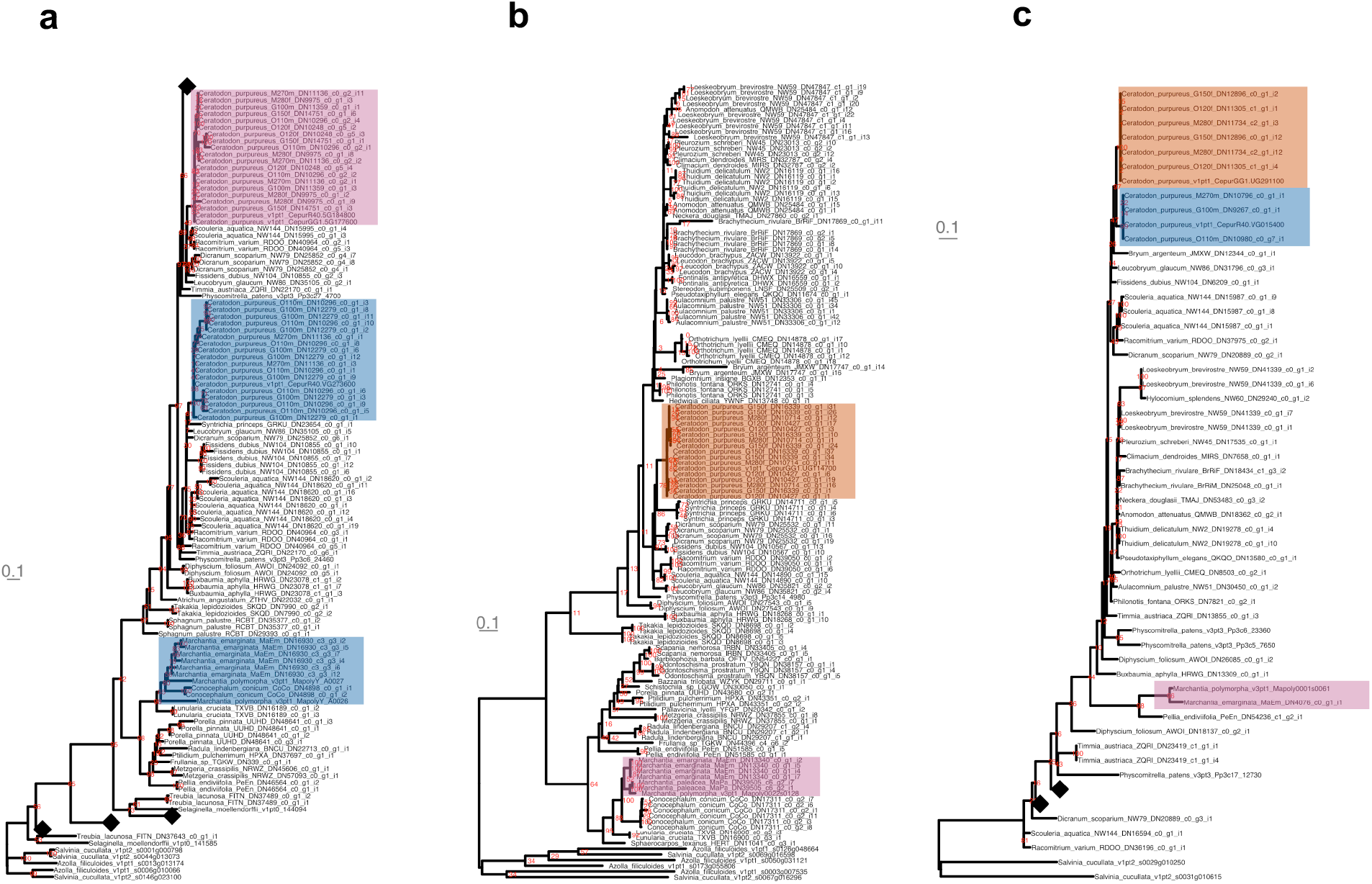
Notable homologous genes in *C. purpureus* and *M. polymorpha*. Trees were built using Maximum-likelihood and bootstrap support is shown in red. Trees were rooted using ferns (*Azolla* or *Salvinia*) as the outgroup. Black diamonds represent collapsed clades and the branch-length scale is in gray. The clades highlighted in blue are V-linked (male), orange are U-linked (female) and purple are noteworthy autosomal homologs. **a**, An *ABC1* gene that is sex-linked in males of *C. purpureus* and *M. polymorpha*. The topology suggests this gene was independently captured between the two lineages. **b**,An *RWP-RK* gene that is sex-linked in females of *C. purpureus* and orthologous to *Marchantia MpRKD*. **c**, Gene tree of sex-linked genes in *C. purpureus* showing they are orthologous to *Marchantia MpFGMYB*.

**Extended Data Table 1.** Sex-biased gene expression patterns in *C. purpureus* for protonemal and gametophore stages. Autosomal genes presented are significantly differentially expressed at an adjusted P≤0.05. U and V-linked genes given are sex-specific (i.e., no evidence of a paralog on the other sex chromosome). Many homologous sex-linked genes are also expressed, however, it is not possible to statistically test for differences in expression between these because they are mapped to different chromosomes. Mean count (MC) Fold change (FC)

|  | Protonema |  | Gametophore |  |
| --- | --- | --- | --- | --- |
| | MC≥1 | FC≥2; MC≥1 | MC≥1 | FC≥2; MC≥1 |
| Male autosomal | 2048 | 317 | 1307 | 256 |
| Female autosomal | 1900 | 366 | 1420 | 264 |
| U | 1006 | - | 1061 | - |
| V | 978 | - | 1054 | - |

Mean count (MC)
Fold change (FC)

## Notes

### Competing Interest Statement

The authors have declared no competing interest.

