## Supplementary Figures and Methods for "The Ceratodon purpureus genome uncovers structurally complex, gene rich sex chromosomes"

**This file includes:**

Supplementary Figures 1-2

Supplementary Methods

### Supplementary Figures

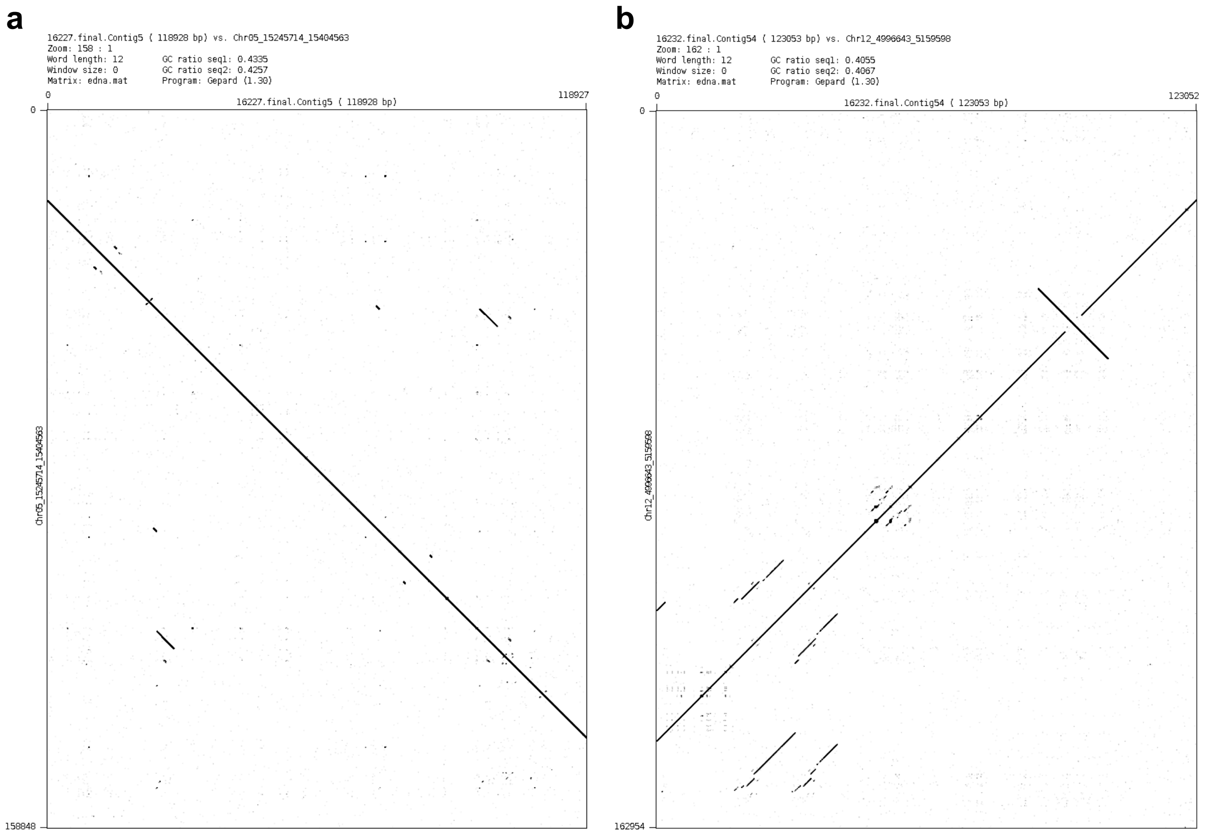


**Supplementary Fig. 1. a,** Dot plot of BAC clone 16227 on a region of Chr_05. This alignment is representative of the high quality BAC clone alignments in 14 of the 18 available BAC clones. **b,** Dot plot of BAC clone 16232 on a region of Chr_12, which is representative of the 3 clones that landed in regions containing a tandem repeat of the genome.

**a**

**b**

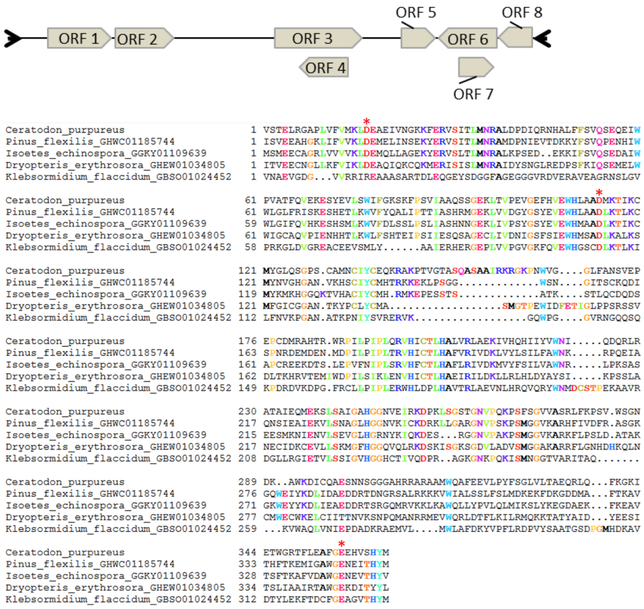


**c**

| Search type | ORF 1 | ORF 2 | ORF 3 | ORF 4 | ORF 5 | ORF 6 | ORF 7 | ORF 8 |
|---|---|---|---|---|---|---|---|---|
| CDD | pfam03962 Mnd1 family | - | PTZ00121 MAEBL | - | - | - | - | - |
| BLAST | HP (Enterobacter cloacae complex) | HP (Klebsormidium nitens) | - | - | - | HP (Enterobacter cloacae complex) | - | - |
| PSI-BLAST | HP (Enterobacter cloacae complex) | HP (Klebsormidium nitens) | - | - | - | HP (Enterobacter cloacae complex) | - | - |
| Hhpred | - | RAG1L, RAG2L/DNA Complex | - | - | - | - | - | - |
| Repbase | - | - | - | - | - | - | - | - |

**Supplementary Fig. 2. Breakdown of the Lanisha transposable element. a,** Schematic representation of a reference Lanisha element. The connecting black line represents a total length of 10,862 bp. Black arrows symbolize terminal inverted repeats and arrowed boxes indicate predicted open reading frames**. b,** Lanisha are found across several plant taxa. Caption of a multiple sequence alignment of putative Lanisha transposase proteins found in the *C. purpureus* genome and in transcriptomes from *Pinus flexilis* (Gymnosperm), *Isoetes echinospora* (Lycopod), *Dryopteris erythrosora* (Fern) and *Klebsormidium flaccidum* (*Klebsormidiophyceae*, previously named *K. nitens*). Sequence identifier of each transcript is indicated. Red stars on top of alignment columns indicate the putative DDE catalytic site. **c,** Summary of homology searches with the eight ORFs predicted in Lanisha reference element. For all searches, E-value cutoff was set at $1x10^{-3}$. Only the best hit is reported. HP: hypothetical protein.

### Supplementary Methods

**Protonemal tissue generation, DNA extraction, and library prep.** All *C. purpureus* protonema (juvenile stage), for all conditions followed the same method for tissue generation unless otherwise noted. For the genome isolates, GG1 and R40, we first blended protonemal tissue from young, less-than-ten day old sub-cultures with a PowerGen 125 (Fisher Scientific, Hampton, New Hampshire, USA) and homogenizers (OMNI-INC 3_750; Omni International, Kennesaw, Georgia, USA) in sterile $dH_2O$ and 1.5mL of moss homogenate was plated on six BCDA plates with cellophane per isolate. This tissue was grown in 16 hours of light at 25°C and eight hours of dark at 21°C until sufficient tissue developed for DNA extraction. For PacBio and Hi-C sequencing tissue was flash frozen in liquid nitrogen for high-molecular weight DNA extraction.

For the PNWB and BUHZ Illumina libraries, genomic DNA was extracted using the Nucleon PhytoPure Genomic DNA Extraction Kit (GE Healthcare) from tissue that was homogenized in liquid nitrogen using a mortar and pestle. Libraries were built using Illumina Regular Fragment, 500bp - 100 ng of Genomic DNA was sheared to 500 bp using the Covaris LE220 and size selected with SPRI using TotalPure NGS beads (Omega Bio-tek) The fragments were treated with end-repair, A-tailing, and ligation of Illumina compatible adapters (IDT, Inc) using the KAPA-HyperPrep kit (KAPA biosystems). The prepared libraries were quantified using KAPA Biosystems next-generation sequencing library qPCR kit and run on a Roche LightCycler 480 real-time PCR instrument. The quantified library was then prepared for sequencing on the Illumina HiSeq sequencing platform utilizing a TruSeq paired-end cluster kit, v3, and Illumina’s cBot instrument to generate a clustered flowcell for sequencing. Sequencing of the flowcell was performed on the Illumina HiSeq2000 sequencer using a TruSeq SBS sequencing kit 200 cycles, v3, following a 2x150 indexed run recipe.

For genomic DNA extraction for PacBio sequencing (libraries PANS and PAPR), intact nuclei were prepared by following the nuclei isolation methods of [114] by grinding approximately 10 grams of biomass in liquid nitrogen and immediately transferring to 200mL of Nuclei Isolation Buffer (NIB) supplemented with 100mM EDTA, pH 8.0. The solution was gently mixed for 10 minutes and filtered with 2 layers of miracloth. Nuclei were pelleted by centrifugation in a swing-bucket rotor at 3,000 rpm for 15 minutes at 4C, washed with NIB 2 times and finally resuspended in 1mL of NIB and an equal volume of 1.5% low-melt agarose. Nuclei plugs were deproteinized and washed according to the methods of Luo and Wing 2003. Intact genomic DNA was retrieved from the agarose plugs by electroelution (BioRad Model 422) at 100 volts for 4 hours in 1X Tris-Acetate-EDTA (TAE), pH 8.3. The resulting genomic DNA was size-verified by pulsed-field gel electrophoresis (PFGE). Long-read sequencing was performed on a PacBio Sequel instrument. Libraries were constructed using SMRTbell Template Prep Kit 1.0 and sized on a SAGE Blue Pippin instrument 10-50kb. Sequencing was performed using a 10-hour movie time and data was generated from a total of 8 SMRT cells per genome.

For Dovetail Hi-C sequencing (libraries IPHH and IPHI), Plant leaf tissue was harvested, flash frozen in liquid nitrogen and kept frozen at -80. Hi-C libraries were constructed at Dovetail Genomics (https://dovetailgenomics.com) using the Dovetail Hi-C kit. Libraries were sequenced using Illumina NovaSeq 6000 Instrument and HiSeq 2500 Instruments, PE150.

For the Bacterial Artificial Chromosomes (BACs) we followed the protocol found in [114] with a few modifications: we supplemented the Nuclei Isolation Buffer (NBI) with 200mM EDTA-pH 8.0 and used *Hin*dIII as the restriction enzyme to partially digest the genome. BAC DNA was isolated from a single bacterial colony and purified on a Qiagen MaxiPrep column. DNA was sheared to 3-4kb using Adaptive Focused Acoustics technology (Covaris, Woburn, MA, USA) and cloned into the plasmid vector pIK96 as previously described [16]. Universal primers and BigDye Terminator Chemistry (Applied Biosystems) were used for Sanger sequencing randomly selected plasmid subclones to a depth of 10x. The Phred/Phrap/Consed suite of programs were then used for assembling and editing the sequence [115–117]. After manual inspection of the assembled sequences, finishing was performed both by resequencing plasmid subclones and by walking on plasmid subclones or the BAC clone using custom primers. All finishing reactions were performed using dGTP BigDye Terminator Chemistry (Applied Biosystems). Finished clones contain no gaps and are estimated to contain less than one error per 10,000 bp.

**Tissue conditions for RNA.** *Ceratodon purpureus protonema:* Petri dishes of homogenized moss tissue were haphazardly chosen for which tray and shelf location to be placed within the growth chamber and rotated daily. As some residual gametophore tissue (if gametophores have already developed on subcultures) occasionally survives blending, we repeated this blending process a second time after six days of growth. We flash froze tissue in liquid nitrogen after the second round of six days of growth.

*Protonemal gene expression and co-expression:* For the gene expression and co-expression lines (see Supplementary Table 5 for isolates and libraries), Petri dishes were haphazardly chosen to be on one of three trays on the same shelf within the growth chamber. Daily the plates were haphazardly rotated within these trays and the trays moved position on the shelf. To capture biological variation for every isolate (i.e., biological replicates), each of the six plates of protonema was frozen in liquid nitrogen separately. RNA was extracted from three of the six biological replicates.

*Modified conditions:* To illicit additional gene expression for use in the genome annotation, we subjected protonemal tissue of the genome isolates, GG1 and R40, to 17 different different conditions. The conditions are described below, which follow the *Physcomitrella patens* Gene Atlas [118], as closely as possible. However, we note these samples were not rotated within the growth chambers. For these conditions, we first homogenized and grew tissue on BCDA for seven days and then subjected the tissue to the new condition for seven additional days, unless otherwise noted, and tissue was flash frozen in liquid nitrogen.

*Agar modifications:* Homogenized tissue was plated on BCD media (i.e., BCDA excluding additional ammonium tartrate [119], BCDA media with 0.5% sucrose, and 0.5% glucose.

*Light treatments:* For the no-light treatment, homogenized tissue was plated on BCDA media supplemented with 0.5% sucrose and the plates covered in aluminum foil, and placed in the growth chamber. For the light treatments, homogenized tissue was plated on BCDA media supplemented with 0.5% glucose and placed in a light box either emitting far-red light (735 nm) and intensity of ~36 $\mu Mols^{-2}s^{-1}$, red light (670 nm) and intensity of ~50 $\mu Mols^{-2}s^{-1}$, yellow light (600 nm) and intensity of ~50 $\mu Mols^{-2}s^{-1}$, blue light (450 nm) and intensity of ~50 $\mu Mols^{-2}s^{-1}$, or green light (530 nm) and intensity of ~50 $\mu Mols^{-2}s^{-1}$ with 18 hours of light and 6 hours of dark at 25°C.

*Hormone treatments:* Homogenized tissue was grown on cellophaned BCDA media for seven days, however, rather than re-homogenize the tissue, it was transferred to BCDA containing 5.0 mg/L of Auxin (1-Naphthaleneacetic acid), 1.0 mg/L of Abscisic acid (both + and – isomers), or 1.0 mg/L of Cytokinin (6-Benzylaminopurine) and grown for an additional seven days.

*External treatments:* Cellophanes containing tissue were transferred to an incubator at 37°C (hot) or a refrigerator at 4°C (cold) for one hour. For dehydration and rehydration, tissue was transferred to a dessicator containing silica gel dessicant and grown for an additional seven days. For rehydration, tissue was transferred to a Petri dish with sterile water for five minutes to rehydrate tissue and returned to the growth chamber for two hours.

*Ceratodon purpureus gametophore gene expression and co-expression:* To generate gametophore tissue (mature stage) to compare gene expression and co-expression, we used the same isolates as protonema (see Supplementary Table 5). Each moss isolate was clonally propagated 16 times each in two magenta jars containing BCDA medium and grown for 122 days with 12 hours of light at 21°C and 12 hours of dark at 10°C. The intention with providing a different temperature for the gametophore stage was to encourage sex expression, however, to our knowledge, sex expression did not occur. Magenta jars were haphazardly chosen to be on one of three trays on the same shelf within the growth chamber and were haphazardly rotated within these trays and the trays moved position within the chamber daily. To capture biological variation for every isolate (i.e., biological replicates), from each of the two magenta jars three distinct patches of gametophore tissue was frozen in liquid nitrogen separately. As a note, some protonemal tissue was still visible in the gametophore jars. Much caution was taken to try and exclude as much protonema as possible without altering gene expression within the gametophore samples, however, we cannot confidently conclude that no protonemal tissue made it in the gametophore samples. One jar of isolate M270m became fungally contaminated after several weeks of growth. Tissue from this jar was collected, but not from the region that had the contamination. Tissue was flash frozen in liquid nitrogen and RNA was extracted from three of the six replicated tissue packets for gametophore tissues.

*GG1 and R40 gametophore:* Homogenized moss tissue was plated on Turface and grown until sufficient gametophore stage tissue developed. Gametophore tissue was harvested by cutting above the Turface surface and flash frozen in liquid nitrogen.

*Ceratodon purpureus sporophyte*: To generate sporophyte tissue (diploid, embryo stage), the G150f and G100m isolates were initially grown on BCDA plates in 18 hours of light at 21°C and six hours of dark at 10°C. Once enough tissue had accumulated, equal amounts of protonemal tissue was blended in 40 mL of sterile d$H_2O$. One mL was plated on a soil and sand amendment (three parts sand and two parts soil with 40 mL of Turface at the bottom) of each of the 2” x 2” pots. The moss suspension for each isolate was placed at equal distances apart to avoid competition for vegetative growth. Pots were randomly placed in trays where position in the tray and trays were rotated weekly. Crosses were grown in a greenhouse in Portland, OR, USA starting in the Spring of 2018 and harvested in the Fall of 2018, at which point the sporophytes reached the elongated and pre-meiotic stage of sporophyte development, and flash frozen in liquid nitrogen.

*Brachythecium rivulare gametophore*: Fertile moss samples with mature female and male gametangia were collected from natural populations of *Brachythecium rivulare* (Brachytheciaceae) and sex determination of the samples was performed. The samples were stored in -80°C until RNA extraction. Two individuals of each sex were pooled to take into account intraspecific variation.

**RNA extraction, library prep, and sequencing**. *Ceratodon purpureus RNA extraction*: The *C. purpureus* protonema and gametophore RNA was extracted using a Qiagen RNeasy Plant Mini Kit (Qiagen, Hilden, Germany) following standard protocols. The *C. purpureus* sporophyte RNA was extracted using an RNeasy Micro kit (Qiagen, Hilden, Germany) following standard protocols. For the sporophyte RNA, we used 25 whole sporophytes, including the capsule and seta, with gametophores detached for each extraction.

The RNA quantity, quality, library prep, and sequencing was done at the Department of Energy (DOE) Joint Genome Institute (JGI) in Walnut Creek, California and the HudsonAlpha Institute in Huntsville, Alabama.

*Protonema library prep and sequencing:* For libraries ATGZB, ATGZG, ATGZP, ATGZO, ATGZH, ATGZN, ATGYY, ATGYZ, ATGZU, ATGYO, ATGZS, ATGZT, ATGYU, ATGYT, ATGYX, ATGYW, ATGZC, ATGZA, ATGYS, ATGYP, plate-based RNA sample prep was performed on the PerkinElmer Sciclone NGS robotic liquid handling system using Illumina's TruSeq Stranded mRNA HT sample prep kit utilizing poly-A selection of mRNA following the protocol outlined by Illumina in their user guide and with the following conditions: total RNA starting material was 1 ug per sample and 8 cycles of PCR was used for library amplification. The prepared library was then quantified using KAPA Biosystems next-generation sequencing library qPCR kit and run on a Roche LightCycler 480 real-time PCR instrument. The quantified library was then then multiplexed with other libraries, and the pool of libraries was then prepared for sequencing on the Illumina HiSeq sequencing platform utilizing a TruSeq paired-end cluster kit, v4, and Illumina's cBot instrument to generate a clustered flow cell for sequencing. Sequencing of the flow cell was performed on the Illumina HiSeq2500 sequencer using HiSeq TruSeq SBS sequencing kits, v4, following a 2x150 indexed run recipe.

*Gametophore library prep and sequencing:* For libraries COZZX, COZZY, COZZZ, CPAAA, CPAAG, CPAAN, CPAAO, CPAAT, CPAAS, CPAAB, CPAAC, CPAAP, CPAAU, plate-based RNA sample prep was performed on the PerkinElmer Sciclone NGS robotic liquid handling system using Illumina's TruSeq Stranded mRNA HT sample prep kit utilizing poly-A selection of mRNA following the protocol outlined by Illumina in their user guide, and with the following conditions: total RNA starting material was 1 µg per sample and 8 cycles of PCR was used for library amplification. The prepared library was then quantified using KAPA Biosystems next-generation sequencing library qPCR kit and run on a Roche LightCycler 480 real-time PCR instrument. The quantified library was then then multiplexed with other libraries, and the pool of libraries was then prepared for sequencing on the Illumina NovaSeq sequencing platform using NovaSeq XP v1 reagent kits, S4 flow cell, following a 2x150 indexed run recipe.

For libraries PWXH and PWXA, stranded cDNA libraries were generated using the Illumina Truseq Stranded RNA LT kit. mRNA was purified from 1 ug of total RNA using magnetic beads containing poly-T oligos. mRNA was fragmented and reversed transcribed using random hexamers and SSII (Invitrogen) followed by second strand synthesis. The fragmented cDNA was treated with end-pair, A-tailing, adapter ligation, and 10 cycles of PCR. The prepared libraries were quantified using KAPA Biosystem's next-generation sequencing library qPCR kit and run on a Roche LightCycler 480 real-time PCR instrument. The quantified libraries were then multiplexed with other libraries, and the pool of libraries was then prepared for sequencing on the Illumina HiSeq sequencing platform utilizing a TruSeq paired-end cluster kit, v3, and Illumina's cBot instrument to generate a clustered flow cell for sequencing. Sequencing of the flow cell was performed on the Illumina HiSeq2000 sequencer using a TruSeq SBS sequencing kit, v3, following a 2x150 indexed run recipe.

*Protonema and gametophore gene expression and co-expression library prep and sequencing:* For libraries COZYT, COZYU, COZYW, COZYX, COZYY, COZYZ, COZZP, COZZS, COZZU, COZZW, COZZA, COZZB, COZZC, COZZG, COZZH, COZZN, COZXO, COZXP, COZXS, COZXT, COZXU, COZXW, COZYG, COZYH, COZYN, COZYO, COZYP, COZYS, COZXX, COZXY, COZXZ, COZYA, COZYB, COZYC, COZZO, COZZT, plate-based RNA sample prep was performed on the PerkinElmer Sciclone NGS robotic liquid handling system using Illumina's TruSeq Stranded mRNA HT sample prep kit utilizing poly-A selection of mRNA following the protocol outlined by Illumina in their user guide, and with the following conditions: total RNA starting material was 1 µg per sample and 8 cycles of PCR was used for library amplification. The prepared library was then quantified using KAPA Biosystems next-generation sequencing library qPCR kit and run on a Roche LightCycler 480 real-time PCR instrument. The quantified library was then then multiplexed with other libraries, and the pool of libraries was then prepared for sequencing on the Illumina NovaSeq sequencing platform using NovaSeq XP v1 reagent kits, S4 flow cell, following a 2x150 indexed run recipe.

*Modified protonema conditions and sporophyte library prep and sequencing*: For libraries GNGYY, GNGYX, GNGYT, GNGYW, GNGYU, GNGYH, GNGZB, GNGYS, GNGYZ, GNGXZ, GNGYP, GNGYO, GNGZA, GNGYC, GNGYG, GNGYA, GNGYN, GNHAA, GNGZZ, GNGZW, GNGZY, GNGZX, GNGZO, GNHAG, GNGZU, GNHAB, GNGZC, GNGZT, GNGZS, GNHAC, GNGZH, GNGZN, GNGZG, GNGZP, GNGYB libraries were built using Illumina RNASeq with PolyA Selection. Plate-based RNA sample prep was performed on the PerkinElmer Sciclone NGS robotic liquid handling system using Illumina's TruSeq Stranded mRNA HT sample prep kit utilizing poly-A selection of mRNA following the protocol outlined by Illumina in their user guide, and with the following conditions: total RNA starting material was 1 ug per sample and 8 cycles of PCR was used for library amplification. The prepared libraries were quantified using KAPA Biosystems next-generation sequencing library qPCR kit and run on a Roche LightCycler 480 real-time PCR instrument. Sequencing of the flowcell was performed on the Illumina NovaSeq sequencer using NovaSeq XP V1 reagent kits, S4 flowcell, following a 2x150 indexed run recipe.

*Brachythecium rivulare:* For *B. rivulare* total RNA was extracted using Tri Reagent (Sigma, St Louis, MO) as described in [120]. RNA library preparation and high-throughput sequencing were performed at FiMM (Finnish Institute for Molecular Medicine) using Illumina HiSeq 2500 sequencing platform. One full lane of 100 base paired-end reads for each sample was sequenced to provide sufficient coverage to provide a representative overview of the expression profile. The transcriptome library was not normalized in order to allow for the detection of differences on the gene expression level between males and females.

**Identification and characterization of Lanisha elements.** From the prediction and the annotation of repetitive elements using REPET, we identified a specific consensus sequence producing relatively high coverage in V chromosome compared to other chromosomes. Detailed characterization revealed that this sequence belongs to a potential novel superfamily of cut-and-paste DNA transposons that we named “Lanisha”. Sequences belonging to this superfamily are 10-12 kb in length, contain 35-40 bp terminal inverted repeats (TIRs) and encode 5 to 8 open reading frames (ORFs) (Supplementary Fig. 2). One ORF, which is conserved across Lanisha elements, was found to present significant similarity with RAG1-type transposases using HMM-based homology search with HHpred tool from HH-suite3 [70] (Supplementary Fig. 2. However, it does not share significant homology with sequences from the RepBase [121] database nor with those from a manually established collection of diverse cut-and-paste transposases [122] using BLAST. Additional homology searches against GenBank non-redundant database using BLASTp and PSI-BLAST revealed the presence of homologous putative transposases among *Klebsormidium nitens* (*Klebsormidiophyceae*) predicted proteins. Two additional Lanisha ORFs presented significant similarity with hypothetical proteins of unknown function from species of the *Enterobacter cloacae* complex. In one of them, we detected homology with a domain found in Mnd1 protein which is involved in homologous chromosome pairing and meiotic double-strand break repair [123] (Supplementary Fig. 2). Because they are repeated in the genome, contain TIRs and encode a potential DDE transposase, Lanisha elements are probable cut-and-paste DNA transposons. Since their transposase is distantly related to those of known transposable elements, Lanisha is proposed to represent a novel superfamily. Finally, using tBLASTn against the GenBank WGS and TSA databases, sequences that are homologous to Lanisha transposase were also detected in sequences obtained from Ferns, Gymnosperms and Lycopods (Supplementary Fig. 2). The Lanisha element reference for *C. purpureus* has been added to NCBI GenBank under MT647524.
